# Evidence of Antibody Repertoire Functional Convergence through Public Baseline and Shared Response Structures

**DOI:** 10.1101/2020.03.17.993444

**Authors:** Matthew I. J. Raybould, Claire Marks, Aleksandr Kovaltsuk, Alan P. Lewis, Jiye Shi, Charlotte M. Deane

## Abstract

The antibody repertoires of different individuals ought to exhibit significant functional commonality, given that most pathogens trigger a successful immune response in most people. Sequence-based approaches have so far offered little evidence for this phenomenon. For example, a recent study estimated the number of shared (‘public’) antibody clonotypes in circulating baseline repertoires to be around 0.02% across ten unrelated individuals. However, to engage the same epitope, antibodies only require a similar binding site structure and the presence of key paratope interactions, which can occur even when their sequences are dissimilar. Here, we investigate functional convergence in human antibody repertoires by comparing the antibody structures they contain. We first structurally profile baseline antibody diversity (using snapshots from 41 unrelated individuals), predicting all modellable distinct structures within each repertoire. This analysis uncovers a high (much greater than random) degree of structural commonality. For instance, around 3% of distinct structures are common to the ten most diverse individual samples (‘Public Baseline’ structures). Our approach is the first computational method to provide support for the long-assumed levels of baseline repertoire functional commonality. We then apply the same structural profiling approach to repertoire snapshots from three individuals before and after flu vaccination, detecting a convergent structural drift indicative of recognising similar epitopes (‘Public Response’ structures). Antibody Model Libraries derived from Public Baseline and Public Response structures represent a powerful geometric basis set of low-immunogenicity candidates exploitable for general or target-focused therapeutic antibody screening.

## Introduction

A key component of the human immune system is the antibody/B-cell receptor (BCR) repertoire, a diverse array of immunoglobulins tasked with identifying pathogens and initiating the adaptive immune response. Broad pathogenic recognition is achieved through enormous variable domain sequence diversity, with an estimated 10^10^ unique heavy variable domains (VH) circulating at any one time from a theoretical set of 10^12^ (or 10^16^-10^18^ full antibodies if light variable domain (VL) combinations are considered (1)).

On antigenic exposure, ‘baseline’ (resting-state) antibodies with sufficiently complementary binding sites to an antigen surface epitope are positively selected. The corresponding parent B cells subsequently migrate to the marginal zone of the lymph nodes, where intentional mutations are introduced to their sequence and only the highest-affinity binders survive in the competition for cognate T-helper cells (2).

Therefore, sequencing antibody repertoires before and during an immune response (e.g. vaccination) can reveal how different people respond to the same antigenic challenge, and can both improve our understanding of immunology and inform future vaccine or therapeutic design (3–5). Similarly, comparing the repertoires of healthy individuals against immunosuppressed (*e.g*. HIV) patients may also make known the origins of increased disease susceptibility (6–8).

However, sequencing an entire antibody repertoire is challenging; they are so large that conventional sequencing techniques, such as Sanger sequencing, do not capture enough of the diversity to be informative. Instead, high-throughput immunoglobulin gene sequencing (Ig-seq) technologies (e.g. Illumina MiSeq) are used. These methods create snapshots that are typically on the order of 10^6^-10^7^ VH and/or VL (unpaired) chains, up to a recent upper bound of around 10^9^ (1, 9, 10). Single-cell sequencing methods, capable of preserving VH-VL chain pairings, are now emerging, however their current throughput yields datasets that are too small to study entire repertoire diversity (11–13).

Since most publicly-available Ig-seq data covers only the VH domain, the vast majority of whole-repertoire analysis has been performed over this region alone. The primary analytical method is currently ‘clonotyping’ (14–16). Clonotyping is a computational technique used to sort sequencing datasets into sets of functionally similar chains based on sequence features, and can be performed in several ways. The most common implementation groups sequences with the same predicted V and J gene transcript origins and above a certain percentage (same length) Complementarity-Determining Region H3 (CDRH3) sequence identity.

Such sequence-based approaches have contributed significantly to our knowledge of core immunology. For example, to estimate the true level of sequence similarity that exists across individuals, Briney *et al*. performed deep sequencing and clonotyping of the circulating baseline VH repertoires of ten volunteers (1). They found that just 0.022% of observed clonotypes were ‘public’ (seen in everyone). In a complementary approach, Greiff *et al*. trained a Support Vector Machine on public and private clonal sequences to identify their high-dimensional features, proving that they have distinct immunogenomic properties (17).

Clonotyping can also be used to detect antigen-specific immunoglobulins, through the identification of expanded clones after vaccination, or those present in unusually high proportions in individuals immune to certain diseases. Explorations of expanded lineages have yielded high-affinity antibodies and T cells against numerous pharmacologically interesting antigens, such as HIV proteins (6), cluster of differentiation proteins (18), botulinum neurotoxin serotype A (19), proteins implicated in type-1 diabetes (20), and many more.

However, clonotyping is only likely to identify a small subset of the true number of functionally equivalent antibodies. This is because it assumes that antibodies require a similar genetic background and high CDRH3 sequence identity to achieve complementarity to the same epitope. In reality, similar binding site structures and paratopes can be achieved from different genetic origins (21) and with surprisingly low CDRH3 sequence identity (22). It is also the case that not every epitope is naturally suited to CDRH3-dominated binding, instead preferring broader engagement by multiple CDRs (23). It is difficult to capture these functionally equivalent antibodies by sequence alone. An alternative approach would be to compare the three-dimensional structures of the antibodies, as binders to a given epitope are likely to adopt a similar geometry with residues capable of recapitulating key binding interactions at equivalent topological locations.

Experimental structure determination (e.g. by X-ray crystallography) remains too slow to solve representative portions of antibody repertoires (24). However, structural annotation approaches are now fast enough to geometrically characterise the individual CDRs of millions of sequences a day with increasing accuracy (25, 26). So far, these analyses have focussed solely on the VH chain, and none have considered the impact of VL on binding site configuration. This can most accurately be captured through variable domain (Fv) modelling, and recent developments have afforded homology approaches with sufficient throughput to analyse meaningful portions of the repertoire (27, 28). For example, a recent prototype structural profiling method that creates representative Fv model libraries from large repertoire snapshots, with applications in developability issue prediction (29).

In this paper, we further refine this repertoire structural profiler, and apply the optimised pipeline to the task of repertoire functional screening. We first analyse 41 baseline antibody repertoires from unrelated individuals, and find that the same representative (‘distinct’) binding site structures are predicted to appear across many individuals (‘Public Baseline’ structures). We also show, through the construction of ‘Random Repertoires’, that this level of structural sharing is far greater than would be expected by chance. Our data therefore represents the first computational evidence that sizeable functional commonality could exist in the baseline repertoires of different people. We then implement the same pipeline on pre- and post-vaccination datasets from three unrelated individuals, detecting a significant increase in structural commonality, and identifying all convergent response structures that may recognise similar epitopes (‘Public Response’ structures). We built Antibody Model Libraries (AMLs) by homology modelling a VH-VL sequence pairing predicted to adopt each Public Baseline or Public Response structure. *In silico* analysis of these AMLs shows that they represent a powerful geometric basis set of low-immunogenicity candidates exploitable for general or target-focused therapeutic antibody screening.

## Results

This study comprises two main investigations. First, we use data from an immunoglobulin gene sequencing (Ig-seq) study by Gidoni *et al*. (30) to investigate the degree of structural overlap in the circulating baseline repertoires of many unrelated individuals. We then use data from a longitudinal Ig-seq flu vaccination study by Gupta *et al*. (5) to measure three individuals’ structural responses to exposure to a common antigen. Both translated Ig-seq datasets were downloaded from the Observed Antibody Space (OAS) database (9), retaining only the 41 Gidoni volunteers with sufficiently deep reads (see Methods).

We used an updated version of our repertoire structural profiling pipeline (29) for improved accuracy in CDR structure and VH-VL interface orientation prediction (see Methods, SI Fig. 4). Briefly, repertoire structural profiling takes as input an antibody/BCR repertoire snapshot containing heavy (VH) and light (VL) chain reads. It eliminates VH and VL chains for which not every CDR is modellable. All modellable VH and VL chains are then sequence clustered to reduce computational complexity. Surviving cluster centres are then paired together and the resulting Fvs that are likely to be successfully modelled are retained. Finally, predicted modellable Fvs with the same combinations of CDR lengths are structurally clustered based on the orientation and CDR loop templates forecast to be used during homology modelling. Antibody Model Libraries (‘AMLs’) can then be built from these representative Fv sequences.

### Structurally Profiling the Baseline Immune Repertoire

We first investigated the structural diversity present in the 41 selected Gidoni baseline repertoire datasets. Separately, each dataset was fed through our structural profiling pipeline to identify the set of sequence diverse modellable VH and VL domains, then the number of predicted modellable Fvs, and finally the number of distinct structures in each dataset (Table 1, full table available as SI Table 2).

**Table 1.**
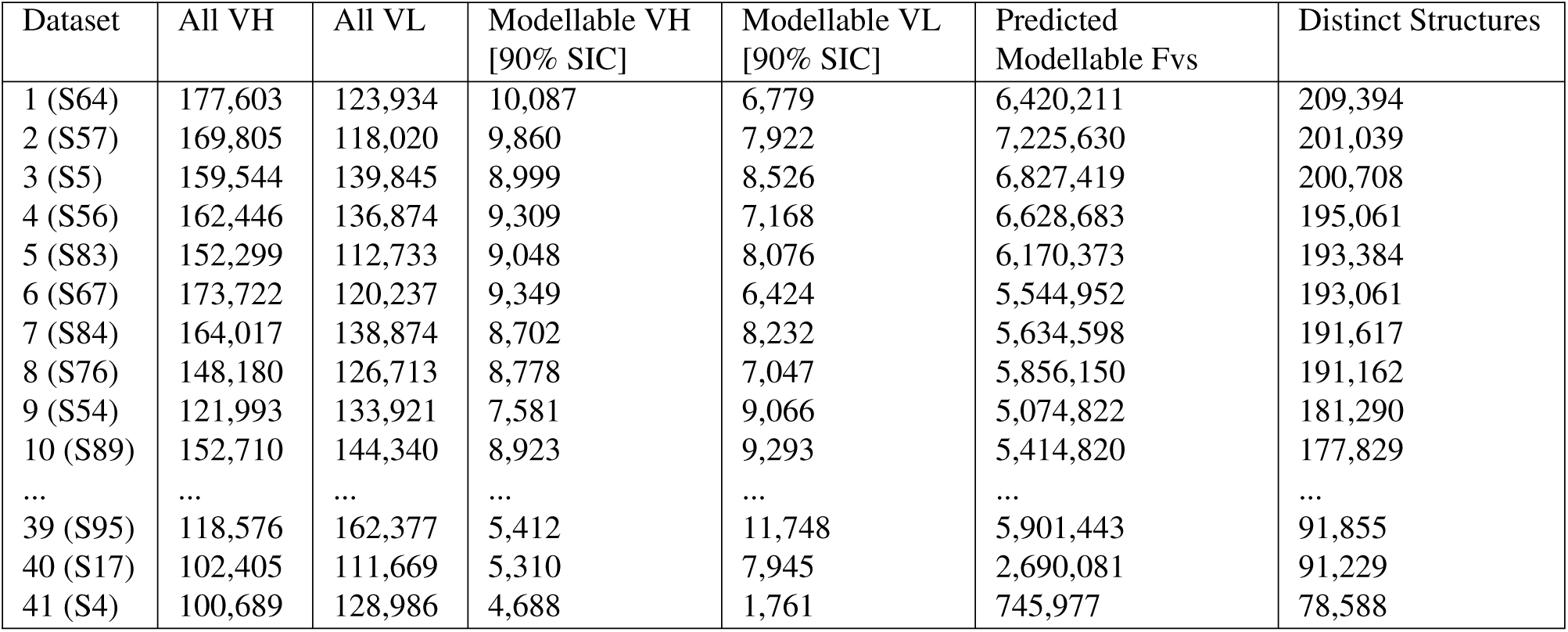
Structurally profiling the baseline repertoire snapshots (30). A full table containing the values for all 41 baseline datasets is available in the Supporting Information (SI Table 2). In order, the columns show: the dataset label, the number of VH and VL reads within each snapshot, the number of FREAD-modellable VH and VL reads (once clustered at 90% sequence identity), the number of predicted modellable Fvs resulting from these VH-VL pairings, and the number of distinct structures (cluster centres) identified in each dataset. SIC = Sequence Identity Clustered.

The most structurally diverse dataset was ‘S64’ (209,394 distinct structures from *∼*6.4M Fvs), and the least was ‘S4’ (78,588 distinct structures, from *∼*750K Fvs). Datasets with a larger number of modellable sequence diverse VHs tended to result in a larger number of distinct structures. Datasets with a moderate/low number of modellable sequence diverse VHs but very large numbers of modellable sequence diverse VLs were amongst the least structurally diverse (*e.g*. ‘S95’). This is consistent with our understanding of both length and structural variability in VH (particularly in CDRH3) relative to VL (31–33).

### Expected Numbers of Distinct Structures (*via*. ‘Random Repertoires’)

To contextualise the numbers of distinct structures observed for each baseline repertoire, we generated ‘Random Repertoires’ to obtain expected numbers of distinct structures assuming each genuine repertoire sampled randomly from modellable, accessible structure space. To achieve this, we derived:

a. The *Modellable Repertoire Structures*: a sample of over 180 million structures built from a random combination of any orientation template, a CDR3 template, and a pair of CDR1/CDR2 templates from the same SAbDab entry (mimicking V gene-encoded predetermination).
b. The *Length-Accessible Repertoire Structures* for each baseline snapshot: the subset of the Modellable Repertoire Structures with a CDR length combination observed in that individual.
c. A *‘Random Repertoire*’ for each baseline snapshot: the appropriate Length-Accessible Repertoire Structures dataset was sampled the same number of times as that individual’s number of predicted modellable Fvs. Clustering these RRs then provided a reference number for the expected number of distinct structures per repertoire, given the depth of sampling in each dataset and assuming random sampling.

To derive a set of Modellable Repertoire Structures, we took the same number of samples as the number of Fvs derived from all baseline repertoire snapshots (183,544,740, SI Table 2). Upon structurally clustering, these samples yielded *∼*28.7M distinct structures over *∼*61.2K distinct combinations of CDR lengths, roughly 100x as many distinct structures as seen in any baseline repertoire sample. However, as each repertoire snapshot typically only contained between 2,000-3,500 different CDR length combinations, many of these 28.7M distinct structures would never be observed in the real data. Therefore, 41 ‘Length-Accessible Repertoire Structures’ datasets were created, limiting the Modellable Repertoire Structures to the CDR length combinations seen in each snapshot. For example, considering only the 3,468 CDR length combinations observed in our most structurally diverse individual (‘S64’) reduced the Modellable Repertoire Structures to a Length-Accessible Repertoire Strucutres dataset of 154.8M structures. This clustered into ~20.8M distinct structures (a 27.5% reduction from the Modellable Repertoire Structures, while the number of CDR length combinations dropped 94%), implying we have good structural sampling over the CDR length combinations typically seen in humans. Every Length-Accessible Repertoire Structures dataset contained a number of randomly-selected structures roughly 20-30 times larger than the number of predicted modellable Fvs observed in the corresponding baseline repertoire.

Finally, 41 separate ‘Random Repertoires’ were created to determine the expected number of distinct structures assuming random structural sampling and given the observed structural sampling depth (see Methods). To do this, each individual’s Length-Accessible Repertoire Structures were sampled randomly, without replacement, the same number of times as the number of predicted modellable Fvs (Table 2).

**Table 2.**
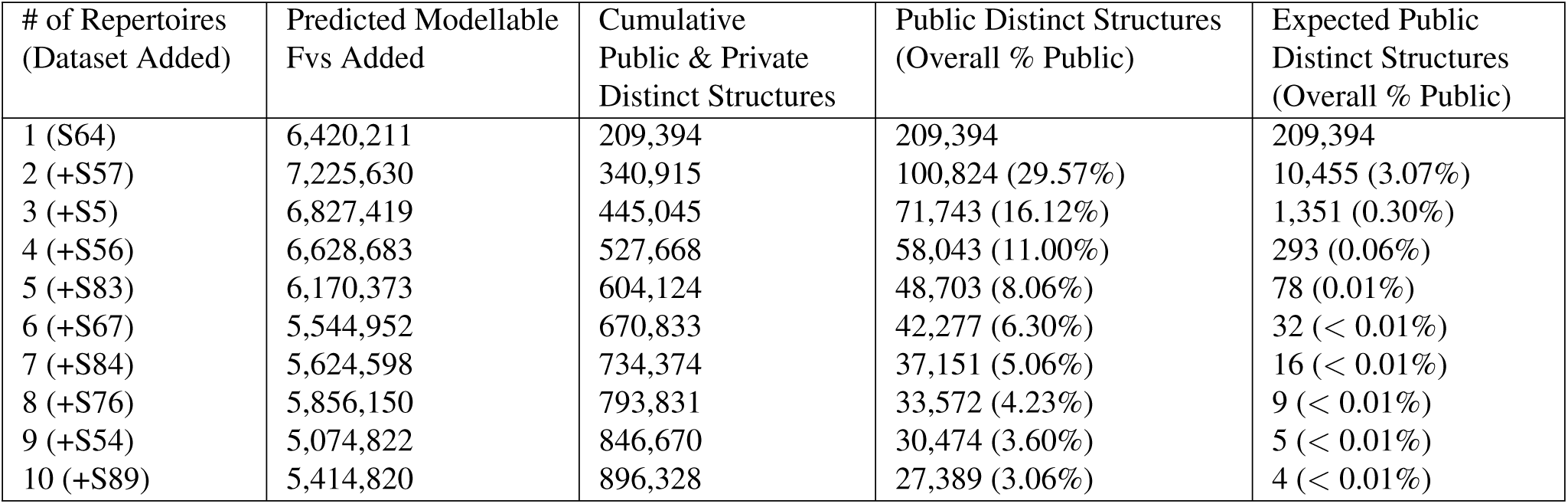
Public structure analysis across the ten most structurally diverse baseline repertoire snapshots. A table tracking the public structures across all datasets is available as SI Table 3. A statistical estimate for the number of public structures was derived by randomly sub-sampling each Random Repertoire to the yield the same number of distinct structures as its equivalent baseline repertoire snapshot. The ‘Public Baseline’ Antibody Model Library was derived from the 27,389 shared structures up to volunteer S89.

| # of Repertoires (Dataset Added) | Predicted Modellable Fvs Added | Cumulative Public & Private Distinct Structures | Public Distinct Structures (Overall % Public) | Expected Public Distinct Structures (Overall % Public) |
| --- | --- | --- | --- | --- |
| 1 (S64) | 6,420,211 | 209,394 | 209,394 | 209,394 |
| 2 (+S57) | 7,225,630 | 340,915 | 100,824 (29.57%) | 10,455 (3.07%) |
| 3 (+S5) | 6,827,419 | 445,045 | 71,743 (16.12%) | 1,351 (0.30%) |
| 4 (+S56) | 6,628,683 | 527,668 | 58,043 (11.00%) | 293 (0.06%) |
| 5 (+S83) | 6,170,373 | 604,124 | 48,703 (8.06%) | 78 (0.01%) |
| 6 (+S67) | 5,544,952 | 670,833 | 42,277 (6.30%) | 32 (< 0.01%) |
| 7 (+S84) | 5,624,598 | 734,374 | 37,151 (5.06%) | 16 (< 0.01%) |
| 8 (+S76) | 5,856,150 | 793,831 | 33,572 (4.23%) | 9 (< 0.01%) |
| 9 (+S54) | 5,074,822 | 846,670 | 30,474 (3.60%) | 5 (< 0.01%) |
| 10 (+S89) | 5,414,820 | 896,328 | 27,389 (3.06%) | 4 (< 0.01%) |

Again taking ‘S64’ as an example, the 6,420,211 samples comprising ‘Random Repertoire S64’ yielded 2,187,257 distinct structures, equating to an average of 2.94 Fvs per distinct structure, compared to 30.66 (10.45x more) Fvs per distinct structure in the genuine repertoire. This provides strong evidence that the modellable portions of antibody repertoires occupy a highly focused region of modellable structure space - roughly 10% of the expected number given the sample size (Fig. 1), and 1% of a theoretical maximum estimate, across the same CDR length combinations.

**Fig. 1.**
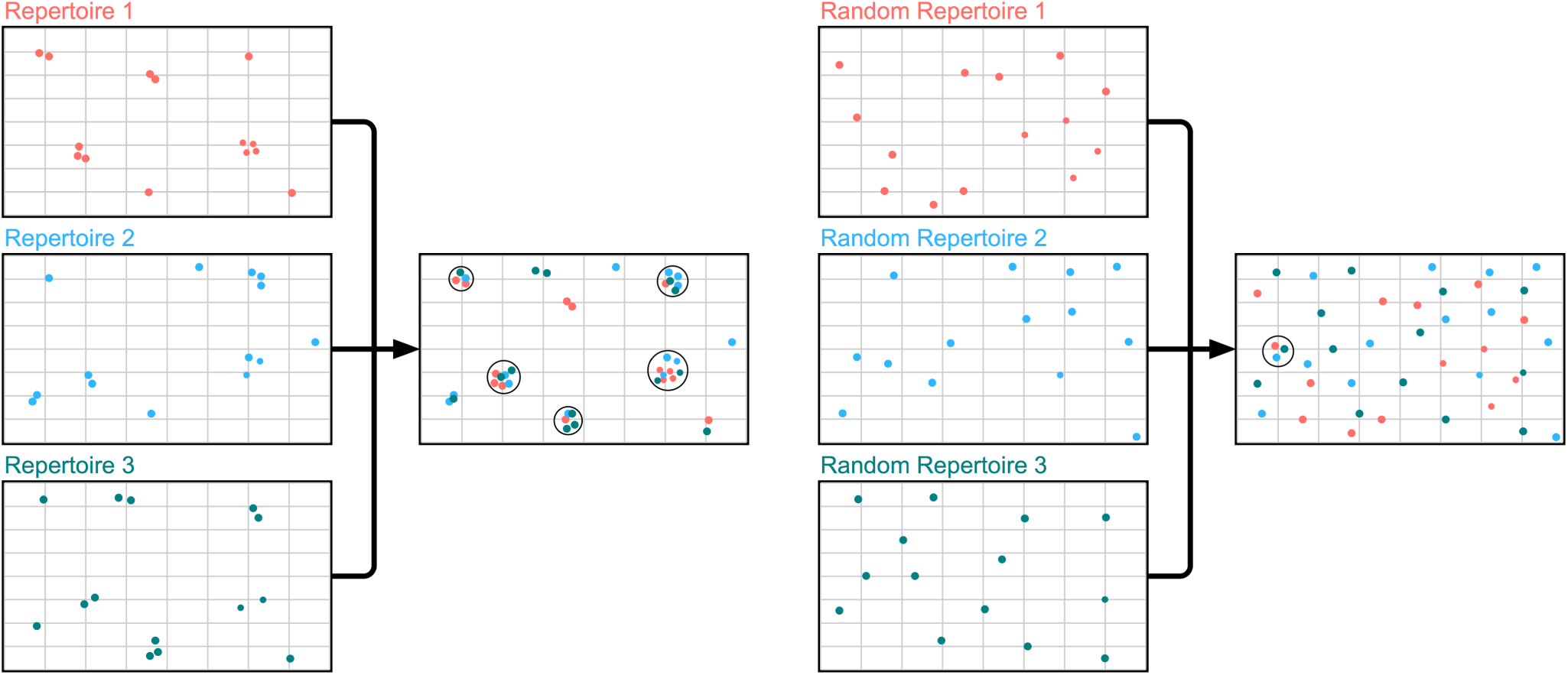
Comparing genuine repertoire snapshot to synthetic ‘Random Repertoires’ (RRs). Each dot represents a distinct structure mapped onto a two-dimensional representation of ‘Length-Accessible Repertoire Structure’ space. The genuine repertoire snapshots of all three individuals (red = repertoire 1, blue = repertoire 2, green = repertoire 3) exhibit focused structural sampling, covering *∼*10% of the space as the corresponding RRs. Overlap analysis shows a high proportion of genuine repertoire distinct structures can characterise an Fv in all three individuals (‘public structures’, represented by black circles). When the same overlap analysis is performed on the equivalent ‘Random Repertoires’, far fewer public structures are observed.

### ‘Public Baseline’ Structures in Unrelated Individuals

We next investigated whether structural commonality exists between baseline repertoire snapshots. This phenomenon would be statistically extremely unlikely by chance, given the focused structural sampling observed in each repertoire. To do this, we performed structural clustering on pairs of repertoire snapshots, looking for evidence of structural overlap (i.e. distinct structures assigned to a predicted modellable Fv seen in both datasets, see Methods and Fig. 2).

**Fig. 2.**
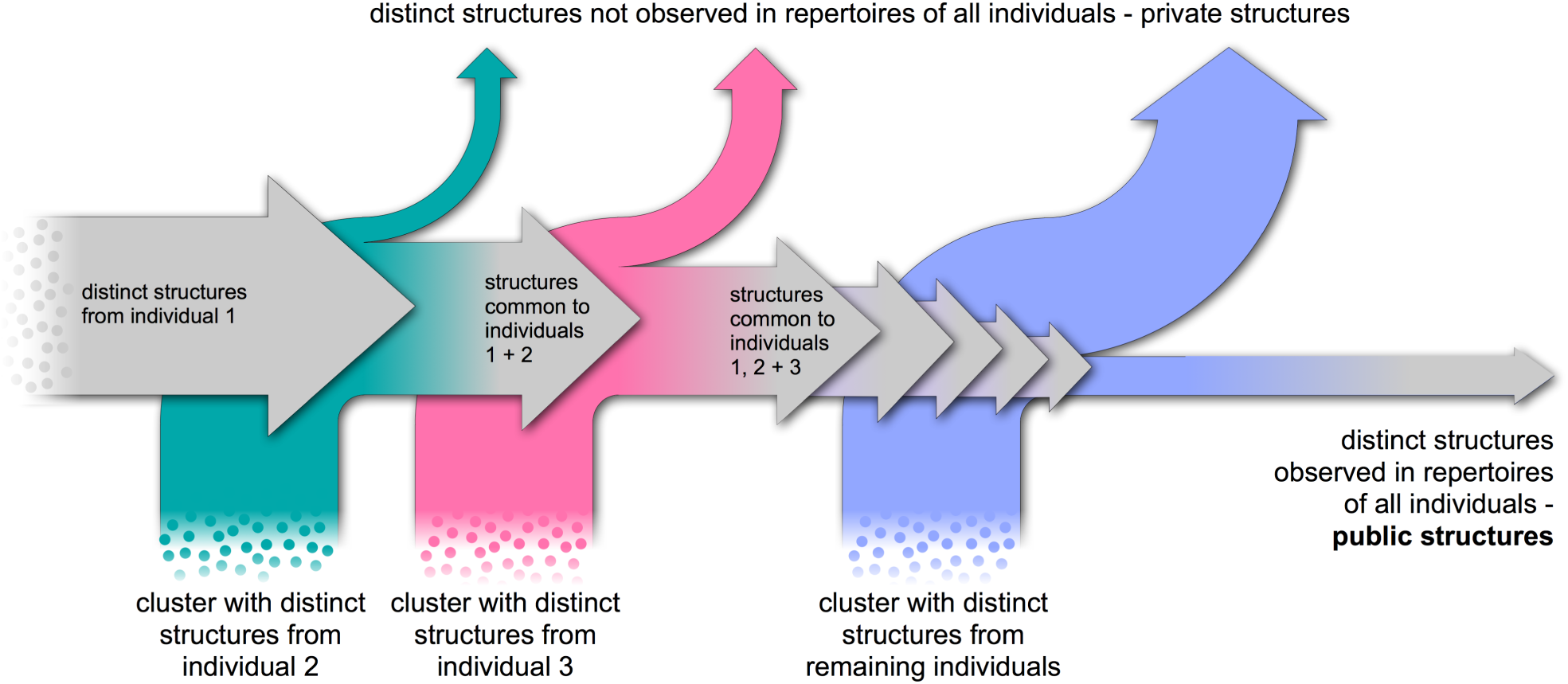
Structural overlap analysis. Datasets are arranged in order of their internal structural diversity (most diverse first). Distinct baseline structures from individual 1 are clustered sequentially with all other repertoire snapshots. Distinct structures present in every tested dataset are classed as ‘public structures’, whereas those that are absent in at least one individual are termed ‘private structures’.

Repertoire snapshots were ordered by their internal structural diversity (‘S64’ first, through to ‘S4’). The 209,394 distinct structures of S64 act as a reference set of cluster centres. The 7,225,630 Fvs from snapshot S57 were then compared to these S64 cluster centres. Structures present in both S57 and S64 were termed public across two individuals, while S64 and S57 distinct structures unique to their own dataset were termed private. Next, the 6,827,419 Fvs from S5 were compared to all public and private distinct structures observed in S64 and S57. We again evaluated the number of public structures, this time present in all three datasets. We repeated this analysis for all remaining baseline repertoire snapshots (first ten results in Table 2, all 41 results in SI Table 3).

To date, all *in silico* analysis of antibody repertoires has suggested that this number should drop rapidly towards 0. For example, a recent clonotype analysis of the baseline circulating repertoire estimated that only around 0.022% of clonotypes were public across ten unrelated individuals (30). However, using our methodology, we found that the number of public distinct structures decreased at a far slower rate, still totalling 27,389 structures after ten unrelated individuals (Table 2). This represents 3.06% of all distinct structures observed up to that point, over 100 times the number of public clonotypes found by Briney *et al*. in their much deeper repertoire samples. Clonotyping our baseline snapshots by the same method, we observed just 26 public clonotypes across the first two repertoires (S64 and S57), and no public clonotypes across the first three repertoires (S64, S57, and S5).

To provide a statistical estimate for how many distinct structures would be expected to be shared across these ten baseline repertoires, the Random Repertoire distinct structures were subsampled to match the corresponding number of baseline repertoire distinct structures (see Methods). In contrast to the genuine repertoires, the Random Repertoires overlapped sparsely, reaching single digits of public structures by just the seventh volunteer (Table 2).

We also tracked the cumulative number of public and private structures over all 41 baseline repertoire snapshots (SI Table 4). Even after the first few most diverse datasets, the deviation from an expected number of distinct structures (given the same ratio of distinct structures:modellable Fvs observed in S64) is quite substantial. This suggests that we might not expect much deviation from our observed fraction of public baseline distinct structures upon deeper repertoire sampling. The existence of such a large number of ‘Public Baseline’ structures would be statistically extremely unlikely without the presence of underlying functional commonality. Clonotyping is fundamentally unable to capture the same depth of signal, even on much deeper sequencing samples, as functional selection is occuring at the level of structural and paratopic similarity, which may not correspond with conservation of gene transcript origin or high CDRH3 sequence identity. Our structural profiling approach is therefore the first computational method to provide supporting evidence for the levels of baseline functional equivalence long hypothesised by immunologists.

### The ‘Public Baseline’ Antibody Model Library

Predicted structures shared between many individuals might represent good starting points for therapeutic development. Their widespread nature could point to their binding versatility, and also to broad immune system tolerance across many individuals, lowering the risk of drug immunogenicity.

We used ABodyBuilder (27) to construct an Antibody Model Library (AML) based on the 27,389 ‘S64’ pairings predicted to adopt a ‘Public Baseline’ structure (as defined by the ten most structurally diverse repertoire snapshots). Some Fvs failed to be entirely homology modelled. For example, occasionally the CDRH3 template clashes irreparably with the CDRL3 template during construction of the full Fv model, necessitating *ab initio* treatment. Overall, 23,700 (86.53%) of 27,389 pairings were entirely homology modelled and comprise our ‘Public Baseline AML’ (SI Dataset 1).

We then investigated the reported species origin of the templates used to model the loops within our ‘Public Baseline AML’ (Table 3). While this is only an approximate measure, as most PDB antibody structures are engineered to some extent, human-origin templates were used considerably more often to model the ‘Public Baseline’ Fvs than would be expected from their abundance in the modelling database (36.47% - 47.07% abundance; 72.63% - 95.69% usage). This further emphasises the value added by starting with human antibody repertoire data, and could indicate that our ‘Public Baseline AML’ structures possess a lower risk of intrinsic immunogenicity to humans.

**Table 3.**
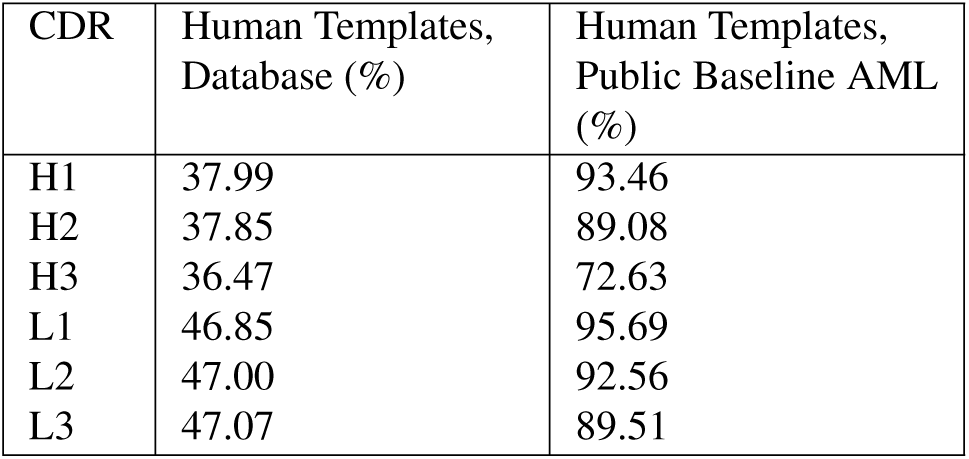
The percentage of each Complementarity-Determining Region’s (CDR’s) templates in the database with reported human origin, against the percentage usage of human-derived templates for CDRs in the ‘Public Baseline’ Antibody Model Library (AML).

| CDR | Human Templates, Database (%) | Human Templates, Public Baseline AML (%) |
| --- | --- | --- |
| H1 | 37.99 | 93.46 |
| H2 | 37.85 | 89.08 |
| H3 | 36.47 | 72.63 |
| L1 | 46.85 | 95.69 |
| L2 | 47.00 | 92.56 |
| L3 | 47.07 | 89.51 |

To test whether our ‘Public Baseline AML’ already contains antibodies proximal to known therapeutics, we mined Thera-SAbDab (34) for all 100% sequence identical structures of WHO-recognised therapeutics, selecting one per therapeutic (see Methods). Of the 66 therapeutics with known structures that had at least one antibody in our ‘Public Baseline AML’ with identical CDR lengths, all had a structural partner in the AML within a C_*α*_ Fv RMSD of 1.84Å, and 37 (56.1%) had a structural partner within 1.00Å Fv RMSD. Eleven therapeutic structures lay within 0.75Å Fv RMSD of a ‘Public Baseline AML’ structure (SI Table 5); these therapeutics spanned a wide range of targets and were primarily successful or promising drugs (4 approved, 5 active in Phase III, 1 active in Phase II, and 2 discontinued).

This result demonstrates that the antibody models within our ‘Public Baseline AML’, without any explicit design, can display high levels of geometric similarity to known therapeutics. Screening libraries based on these commonly shared structures holds significant promise for finding novel low-immunogenicity therapeutics.

### Structurally Profiling a Flu Vaccine Response

Clonotyping is commonly used in antibody drug discovery to identify ‘expanded clones’ - novel genetic lineages present after vaccination/infection but that were absent, or low concentration, beforehand (14). Here, we applied our repertoire characterisation method to investigate whether we could identify an analogous structural response to vaccination.

To this end, we used a longitudinal 2009 seasonal flu vaccination study by Gupta *et al*. (5), in which three unrelated individuals (‘V1-3’) were sequenced at many time-points before and after vaccination. Sequences were again downloaded from the OAS database, yielding ‘Before Vaccination’ and ‘After Vaccination’ datasets for each individual, according to the protocol described in the Methods. Using the same repertoire structural profiling protocol as above, we calculated the number of distinct structures observed in each individual before and after vaccination (SI Table 4).

To obtain an estimate for the degree of structural commonality pre- and post-vaccination, we again used a greedy clustering approach to evaluate the structural overlap between the ‘Before Vaccination’ datasets, and between the ‘After Vaccination’ datasets, separately (Fig. 3a, 3b). The first dataset in each overlap assessment was the most structurally diverse (*i.e*. the ‘V3’ individual before vaccination, and ‘V1’ after vaccination).

**Fig. 3.**
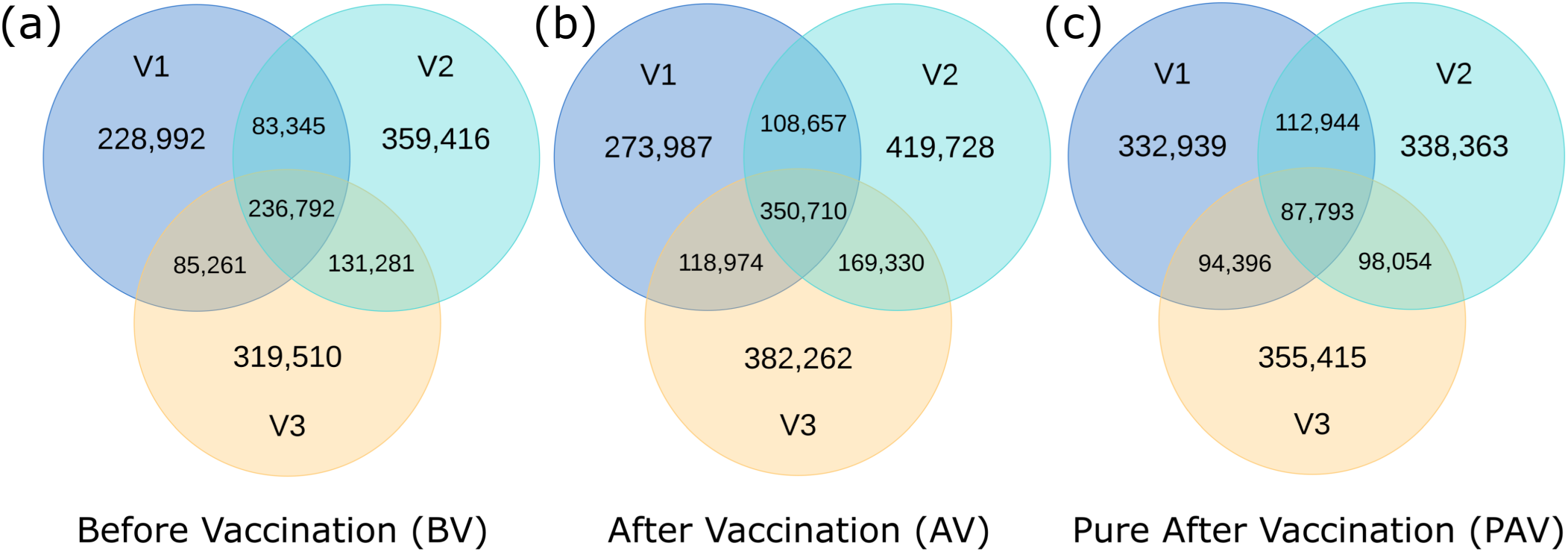
Venn diagrams showing the structural overlap between each individual’s ‘Before Vaccination’ dataset (a), ‘After Vaccination’ dataset (b), and ‘Pure After Vaccination’ dataset ((c), distinct structures arising only after vaccination). Total distinct structures: Before Vaccination - 1,444,597; After Vaccination - 1,823,628; Pure After Vaccination - 1,419,904. V1-V3 = Volunteer 1-3.

Again, a significant number of public distinct structures were observed in ‘V1’,’V2’, and ‘V3’ (‘Public Before Vaccination’ structures, 17.78% (236,792/1,444,597) of all ‘Before Vaccination’ distinct structures). This indicates that the identification of ‘Public Baseline’ structures in the previous section was unlikely due to serendipitous Ig-seq amplification bias. Interestingly, 17.78% is a similar percentage of sharing as that seen after three baseline snapshots (16.12%; 71,743/445,045).

The degree of structural sharing appears to increase after vaccination, with 19.23% (350,710/1,823,648) public structures across the three volunteers. This is consistent with a degree of repertoire structural convergence driven by exposure to the same pathogenic epitopes.

To derive these convergent structures, the structural overlap between each individual’s ‘Before Vaccination’ and ‘After Vaccination’ datasets was measured, only retaining ‘After Vaccination’ pairings that could not be clustered into the same individual’s ‘Before Vaccination’ distinct structures. ‘V1’ remained the most structurally diverse dataset, with 628,072 ‘Pure After Vaccination’ distinct structures. The overlap between these ‘Pure After Vaccination’ pairings (Fig. 3c) was then compared. This yielded a mixed picture of convergent and private vaccination response structures - 27.7% (393,187/1,419,904) of distinct structures were shared with at least one other individual, and 6.18% (87,793/1,419,904) were shared across all three individuals - which we term ‘Public Response’ structures.

There are two potential causes of overlap in the ‘Pure After’ vaccination set. One is a genuine common structural response to vaccination, while the other is that the initial baseline repertoire was under-sampled - *i.e*. the overlap reflects residual shared baseline structures. As a second test for baseline deviation, beyond absence before vaccination, we compared how many of the 27,389 ‘Public Baseline’ distinct structures were within 1Å of a ‘Public Before Vaccination’ binding site, versus the number within 1Å of a ‘Public Response’ Structure binding site. We observed that 80.0% (21,922/27,389) of ‘Public Baseline’ structures were within 1Å of a ‘Public Before Vaccination’ structure, compared to just 24.2% (6,621/27,389) proximal to a ‘Public Response’ structure. This provides further evidence that a proportion of these convergent ‘Public Response’ structures reside in a distinct region of structural space and could harbour epitope-specific binding geometries. We have built a ‘Public Response AML’ based on these 87,793 shared structures, with 74,181 Fvs (84.4%) successfully homology modelled (SI Dataset 2).

## Discussion

In this work, we have structurally profiled antibody repertoires to capture new insights into the baseline and antigen-responding immune system, and to create novel libraries of antibody model structures that could be exploited for immunotherapeutic discovery.

All of the structural analysis in this paper is limited to the antibody chains that are currently predicted to be modellable, and so there remain regions of natural structural space uninvestigated. Nevertheless, we show that antibody repertoires tend only to explore highly focused regions of currently-modellable structural space (*∼*10% of the diversity expected if templates were explored randomly across the same combinations of CDR lengths). This suggests that the current degree of structural commonality would remain across the (as yet) unmodellable regions of structural space.

The enormous sequence diversity exhibited across baseline antibody repertoires has long appeared to run contrary to the theory of baseline functional commonality. Here we have shown that, at least from a structural perspective, there is considerable opportunity for functional commonality across the circulating resting-state repertoires of unrelated individuals (*∼*3% of observed distinct structures are public across 10 individuals). The theoretical chemical diversity that could be displayed on each of these scaffolds is large, so many of these grouped binding sites will not be complementary to the same antigen epitope. However, there is good reason to believe that a certain proportion are, as geometric similarity is a likely prerequisite of functional commonality, and our structural clustering approach offers a route to detecting and analysing these antibodies. This knowledge could then be harnessed in vaccinology - for example, identifying an epitope targetable by a ‘Public Baseline’ structure may lead to a more reliable and convergent response.

We hypothesise that human ‘Public Baseline’ structures are more likely to display low levels of human immunogenicity and be versatile binders. Building full three-dimensional variable domain models of these distinct structures (an Antibody Model Library) produced geometries that were very close to several approved and late-stage active therapeutic antibodies targeting diverse antigens. To chemically elaborate this ‘Public Baseline’ structural basis set, a phage display library on the order of 10^6^-10^7^ sequence-unique human antibodies could be created from the many different Fv sequences predicted to adopt each public distinct structure.

Target-focused screening libraries against immunodominant epitopes are commonly derived through sequence analysis of longitudinal Ig-seq studies that track the immune response of many individuals to the same antigen. We show that when our methodology is applied to a longitudinal flu vaccination case study, we detect a higher level of structural convergence, commensurate with response to similar epitopes on the same antigen. We can also derive a large number of ‘Public Response’ structures, with divergent structural characteristics from the baseline repertoire. These could contain useful binding site structures exploitable for antigen-specific library design.

There are inevitable biases in structurally profiling human antibody repertoire data to suggest antibody leads for drug discovery. One such biased property is CDRH3 length: very short CDRH3 lengths will be under-sampled through their sparsity in natural human sequences (29), while very long CDRH3 lengths will be under-sampled because they are more difficult to homology model accurately. While inherent immunogenicity should be diminished by virtue of using naturally-expressed sequences, other developability issues are still possible, as not every human antibody has the biophysical properties ideal for large-scale manufacture and long-term storage (29).

Nevertheless, we believe that our approach should find immediate applicability in both *in silico* and *in vitro* screening. We have made available the ‘Public Baseline’ and ‘Public Response’ Antibody Model Libraries for further investigation, and will continue to build and share the Antibody Model Libraries derived from other unpaired and paired VH+VL datasets in the Observed Antibody Space database (9).

## Methods

### Immunoglobulin Gene Sequencing Datasets

The cleaned and translated antibody repertoire datasets (5, 30) were downloaded directly from the Observed Antibody Space (OAS) database (9). For the Gidoni data (30), only individuals for whom *>* 100,000 IgM VH and *>*100,000 VL sequences were recorded were analysed. In our analysis of Gupta et al. (5), we used all three individuals (‘V1’ = ‘FV’, ‘V2’ = ‘GMC’, and ‘V3’ = ‘IB’). The ‘Before Vaccination’ data was defined as all VH and VL sequences recorded at 8 days, 2 days and 1 hour before vaccination. The ‘After Vaccination’ data was defined as all VH and VL sequences recorded at 1 week, 2 weeks, 3 weeks, and 4 weeks after vaccination. Sequences recorded 1 hour and 1 day after vaccination were discarded to avoid ambiguity. The ‘Pure After Vaccination’ data contained ‘After Vaccination’ sequences that did not fall into the structural clusters defined by each individual’s ‘Before Vaccination’ repertoires. The seminal work in which ‘FV’, ‘GMC’, and ‘IB’ were vaccinated is detailed in Laserson *et al*. (4), however the data we use derives from Gupta *et al*. (5), who re-analysed each antibody repertoire snapshot with Illumina sequencing.

### Repertoire Structural Profiling Pipeline

The described structural profiling pipeline was optimised from the protocol reported in the Supporting Information of Proc. Natl. Acad. Sci. (2019) 110(6):4025-4030 (29).

#### CDR modellability analysis

Each sequence was first numbered using ANARCI (35) according to the IMGT numbering scheme (36), and the closest framework region (variable domain with North-defined CDRs (31) excised) in the SabDab (23) database (12^th^ February 2019) was identified. In the IMGT numbering scheme, the North CDRs lie between the following residue numbers - CDRH1: 24-40; CDRH2: 55- 66; CDRH3: 105-117; CDRL1: 24-40; CDRL2: 55-69; CDRL3: 105-117.

FREAD (37, 38) was then used to attempt to map each Ig-seq sequence to a length-matched North CDR template. The FREAD CDR databases were timestamped to 12^th^ February 2019, and contained the following numbers of templates - CDRH1: 2,526; CDRH2: 2,575; CDRH3: 2,502; CDRL1: 2,355; CDRL2: 2,373; CDRL3: 2,376. All loop templates contain the North-defined CDR loop and 5 ‘anchor residues’ before and after the loop. Selection of CDRH3 templates was performed according to a bespoke set of Environment-Specific Substitution Score (ESS) thresholds established using Ig-seq data: Lengths 5-8, ESS *≥* 25; Lengths 9-10, ESS *≥* 35; Lengths 11+, ESS *≥* 40 (see SI Methods). Each template surpassing the threshold was subsequently grafted onto the corresponding framework anchor residues. The loop template with the lowest calculated C_*α*_ anchor RMSD was selected. Any sequences for which at least one loop could not be modelled were removed from the dataset.

#### Sequence clustering

The modellable chains were then sequence clustered using CD-HIT (39) at a 90% sequence identity threshold, to reduce the number of VH-VL pairing comparisons to a computationally-tractable number.

#### Predicting modellable VH-VL orientations

The 20 most important VH-VL interface residues for orientation prediction were derived; a sequence identity of 85% over these 20 residues resulted in an orientation RMSD of *≤* 1.5Å *∼* 80% of the time (see SI Methods).

All remaining VH and VL domains after sequence clustering were paired together, and their 20 key interface residues were recorded. If the sequence identity over these residues was *≥* 85% to at least one of 1,129 reference Fvs, the interface was classed as modellable - otherwise the VH-VL pairing was discarded. If multiple reference Fvs shared *≥* 85% identity, the predicted modellable VH-VL pairing inherited the orientation parameters of the Fv reference with highest sequence identity.

#### Identifying distinct structures

At this stage, each predicted modellable VH-VL pairing (Fv) has eight associated parameters: its orientation template, its six CDR templates, and a length vector recording the combination of North CDR lengths (31) present in its binding site. Fvs were then structurally clustered to identify ‘distinct structures’ according to the following process. First, identically-predicted binding sites (for which the eight predicted parameters were the same) were identified. The retained pairing was randomly chosen, except in the overlap studies - if one of the pairings was present as a distinct structure of the first dataset, this pairing was selected and recorded as a shared structure across both repertoires.

Next, singleton length clusters were identified and assigned as separate distinct structures, avoiding inaccurate RMSD comparisons between loops of differing length. The remaining interfaces were split by their CDR length combinations, and were greedily clustered with all other pairings sharing the same length vector as follows:

1. Select the first pairing as a distinct structure (cluster centre).
2. Select the next pairing. If the orientation RMSD to all existing cluster centre orientation templates exceeds 1.5 Å, classify the new pairing as a distinct structure. Otherwise:
3. Calculate the RMSD between the CDR templates of the new pairing with those of all existing cluster centres using the formula:

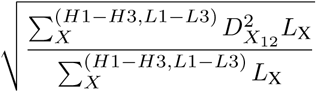

where the sum over X refers to each of the six CDRs, L_X_ is the length of North CDRX, and D_*X*_12 is the C_*α*_ RMSD between the CDRX in Fv 1 and Fv 2. If this value exceeds 1 Å to all existing structural cluster centres, the pairing is assigned as a distinct structure. Otherwise the pairing is stripped from the dataset.
4. Return to step 2 until all pairings have been analysed.

### Overlap Comparison

To identify shared structures between two Ig-seq repertoire snapshots, the distinct structures from the first snapshot were listed followed by all predicted modellable Fvs of the second repertoire snapshot, as an input file to the clustering algorithm. The greedy clustering ensured that all distinct structures from the first dataset remained as cluster centres, and allowed for the identification of pairings in the second dataset that were also predicted to occupy the same structural neighbourhood.

### ‘Random Repertoires’

To contextualise the structural diversity displayed in human antibody repertoires, we derived ‘Random Repertoires’ (RRs) according to the following method. First, a set of Modellable Repertoire Structures (MRS) was generated. When generating a structure, one of any of the 1,129 orientation templates, 2,502 FREAD CDRH3 templates, and 2,376 FREAD CDRL3 templates were available for selection. To mirror the genetics of the immune system (as they would be encoded on the same V gene transcript), CDR1 and CDR2 templates were restricted to being selected from the same SAbDab structure, limiting our choice to one of 2,519 CDRH1/2 templates and 2,345 CDRL1/2 templates. Each of these five sets was randomly sampled over 180 million times to create the MRS dataset. This was then filtered to create 41 Length-Accessible Repertoire Structure (LARS) datasets, containing only the combinations of CDR lengths observed in each baseline repertoire snapshot. Finally, RRs were created by sampling each LARS set the same number of times as the number of predicted modellable Fvs in the corresponding baseline repertoire snapshot.

To obtain statistically expected values for structural overlap across individuals, the distinct structures from ‘RR_S64’ were randomly subsampled the same number of times as the number of distinct structures seen in ‘S64’ itself, yielding random distinct structure samples occupying the same proportion of LARS-space. The process was repeated for each RR dataset, normalising to each respective baseline repertoire snapshot. Overlap comparison was then performed as described above, starting from the ‘RR_S64’ distinct structures, followed by all the pairings that fell into the selected ‘RR_S57’ distinct structures, *etc*.

### Clonotyping

Clonotyping was performed to group antibodies with the same closest V and J gene, and identical CDRH3 sequences, as in Briney *et. al*. (1).

### Antibody Model Library Construction

Antibody model libraries (AMLs) were constructed with a parallel implementation of ABodyBuilder (27), using the FREAD (37, 38) Environment Specific Substitution Scores derived from Ig-seq benchmarking (see CDR Modellability Analysis). Some predicted modellable Fvs are not entirely homology modellable, as loop modellability is considered on a per-chain basis and does not take into account inter-chain loop clashes that become evident only upon full Fv homology modeling. Fvs that required any degree of *ab initio* modelling to fix such issues were trimmed out of the dataset.

### Comparison to Antibody Therapeutics

The set of 89 therapeutics with 100% sequence identical structures (as of November 2019) were retrieved from Thera-SAbDab (34). A single structure was chosen for each therapeutic for the RMSD analysis; if multiple structures were available, we selected unbound structures with the best resolution. RMSD comparisons were only made between therapeutics and AML structures with matching combinations of CDR lengths. Fv RMSD was calculated over all C_*α*_ atoms after alignment of backbone atoms, using an in-house script.

## ACKNOWLEDGEMENTS

This work was supported by an Engineering and Physical Sciences Research Council and Medical Research Council grant [EP/L016044/1] awarded to MIJR, a Biotechnology and Biological Sciences Research Council (BBSRC) grant [BB/M011224/1] awarded to AK, and funding from GlaxoSmithKline plc, UCB Pharma Ltd., AstraZeneca plc, and F. Hoffmann-La Roche AG.

## AUTHOR CONTRIBUTIONS

M.I.J.R., C.M., and C.M.D. designed research; M.I.J.R. performed research; M.I.J.R., C.M., A.K., A.P.L., J.S., and C.M.D. analyzed data; and M.I.J.R., C.M., and C.M.D. wrote the paper.

