## Supporting Information for "Evidence of Antibody Repertoire Functional Convergence through Public Baseline and Shared Response Structures"

Raybould MIJ, Marks C, Kovaltsuk A, *et al.*

Corresponding Author: Charlotte M. Deane


### Methods

**Immunoglobulin Gene Sequencing (Ig-seq) Environment-Specific Substitution Scores.** The original antibody Environment-Specific Substitution Score (ESS) threshold used by FREAD (25 for all loops, and loop lengths) was derived by Choi and Deane (1, 2) based on benchmarking of performance on the Protein Data Bank (PDB). This involved first excising the CDRH3 loop from known PDB antibody structures, and then feeding the framework structure and excised CDRH3 loop sequence into FREAD, asking it to predict the next-best (non-self) PDB CDRH3 template for modelling, and measuring the achieved root-mean-squared deviation (RMSD). However, the CDRH3 loops in Ig-seq samples of natural antibody repertoires deviate significantly from the heavily-engineered PDB, which has high redundancy and many closely-related structures that help to improve mean performance. To show this, we calculated the typical ESS value of the top-ranked FREAD CDRH3 template for an Ig-seq study of natural antibodies (**Fig. 1a**), comparing it to the ESS of the top-ranked FREAD CDRH3 template obtained when modelling-in PDB CDRH3 loops (**Fig. 1b**, blinding access to the same structure as a template). The highest ranked template has the lowest anchor residue $C_\alpha$ RMSD, after surpassing the baseline threshold of ESS 25. The Ig-seq data benefits from far fewer high ESS scores, so application of the original thresholds on Ig-seq data would be expected to achieve below-headline performance. To both estimate our true performance on Ig-seq data, and derive new CDRH3 loop ESS thresholds more appropriate for Ig-seq data, we subsampled the PDB comparison set to match the top-ranked ESS distributions seen for each length bin in the aforementioned Ig-seq sample. Based on this sample, we predict that we should achieve a good accuracy (mean RMSD of 2.54Å) with acceptable coverage on Ig-seq data using the following CDRH3 ESS cutoffs: Lengths 5-8, ESS $\geq$ 25; Lengths 9-10, ESS $\geq$ 35; Lengths 11+, ESS $\geq$ 40.

**Determining Interface Residues Key for Orientation Prediction.** The 1,129 sequence non-redundant Fvs with resolution $\leq$ 2.5 Å were taken from the SAbDab database (3) ($12^{th}$ February 2019), and all residues found to lie in the VH-VL interface were identified. This was achieved by first calculating the relative solvent accessible surface area ($SASA_{rel}$, Shrake-Rupley Algorithm (4)) to determine the absolute SASA of each residue and dividing this number by the theoretical maximum SASA for that residue. The $SASA_{rel}$ of each position in the complex was then compared to the value for the equivalent position in the separated VH and VL chains (coordinates of the partner chain deleted). The resulting 52 residue positions (**SI Table 1**), which appeared in at least 80% of complexes, and whose $SASA_{rel}$ was reduced by an average of at least 5% in at least 10% of those complexes, were taken forward. To reduce this further, we performed a Random Forest regression analysis [standard scikit-learn implementation, 500 estimators] over these 52 residues to the 6 ABangle parameters (5) that have been shown to characterise VH-VL orientation. First we confirmed that the 1129 interfaces constituted a representative sample of all ABangle parameter space - essential to learn genuine residue to ABangle parameter responses. Each interface was then flattened and one-hot-encoded (21 columns for each position, for the 20 natural amino acids or a deletion/missing residue) to yield a 1129x1092 matrix which was separately regressed against each ABangle parameter (6 x (1129x1) vectors). Out-of-bag validation was used to estimate $R^2$ values, and showed predictive performance ranging from an estimated $R^2$ of 0.35-0.54. We calculated feature importance, and derived a new one-hot-encoded interface for each complex using only the 20 positions (**SI Table 1**) that were present in the top-5 most important features across the 6 parameters. Predicted performance dropped by an average estimated $R^2$ of only 0.05 across the six parameters (min: 0, max: -0.11).

The coverage and accuracy of the 52 and 20 residue interface definitions at predicting orientation RMSD was also assessed. Orientation RMSD between two complexes was measured by first aligning their VH domains and measuring the $C_\alpha$ distances between common VL positions, then by aligning their VL domains and measuring the $C_\alpha$ distances between common VH positions, and finally dividing by two. A 'correct/incorrect' orientation threshold RMSD of 1.5 Å was chosen by measuring the variation in pairwise orientation RMSD observed for sequence-identical SAbDab complexes, *i.e.* an estimate for the experimental limitation on what constitutes a 'correct' orientation RMSD. A threshold of 1.5 Å captures 92% of sequence identical Fvs (see **SI Figure 2**). An orientation sequence identity threshold was then chosen for the 20-residue and 52-residue interface definitions that balanced acceptable coverage with a high proportion of Fvs within 1.5 Å above the threshold (see **SI Figure**

DRAFT

3). We chose 82% for the 52-residue definition, and 85% for the 20-residue definition. Coverage was comparable and accuracy only slightly reduced (80.2% to 77.8%) on narrowing the interface definition to 20 residues.

DRAFT

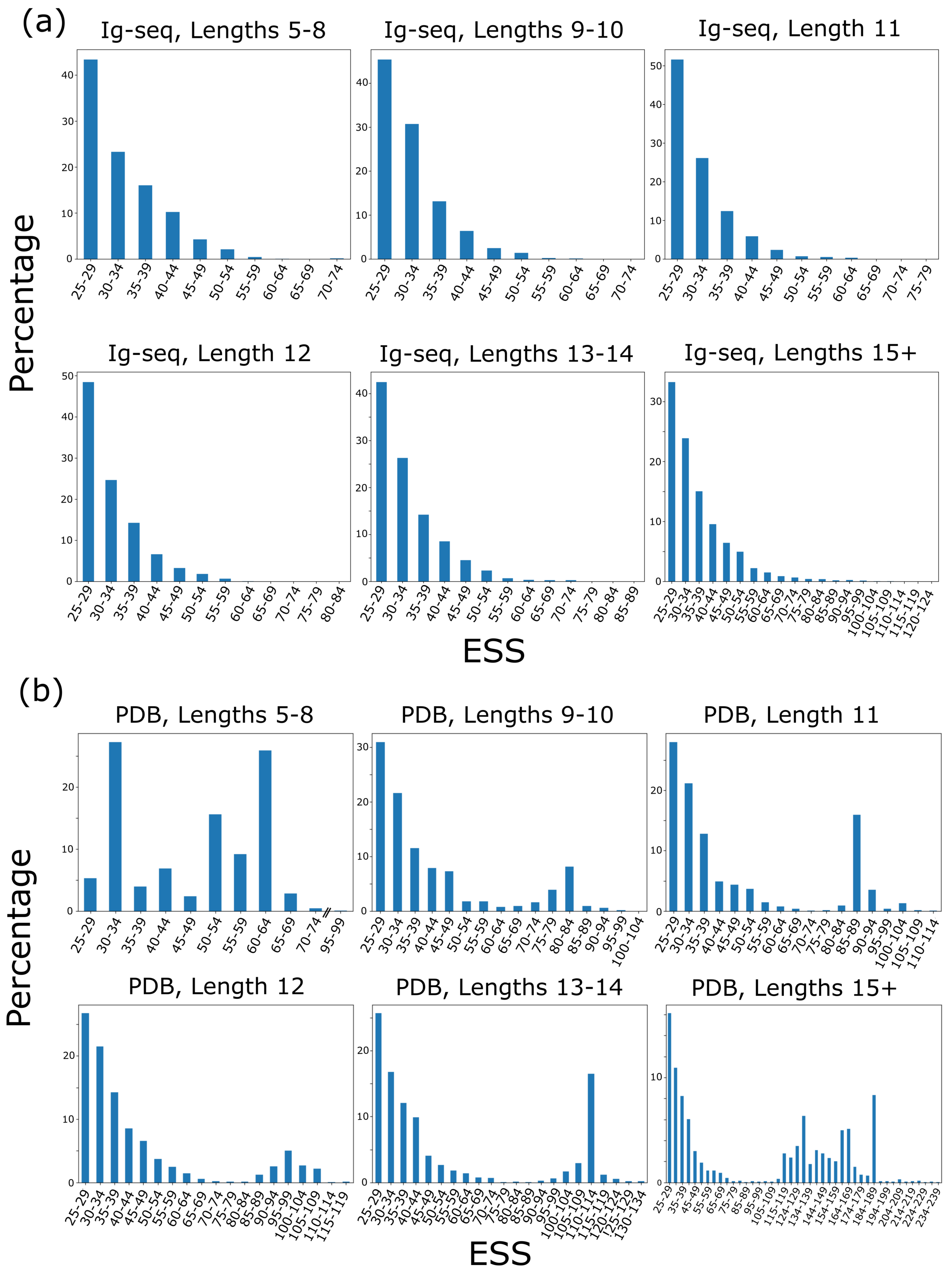


**Fig. 1.** The percentage of each FREAD (1, 2) top-ranked CDRH3 templates with an Environment Specific Substitution Score (ESS) within the labelled bin for (a) a typical Ig-seq dataset, and (b) the Protein Data Bank (blinded to self). The two sets have very different distributions; notably Ig-seq datasets rarely contain CDRH3 loops with extremely high ESS scores to dataset templates.

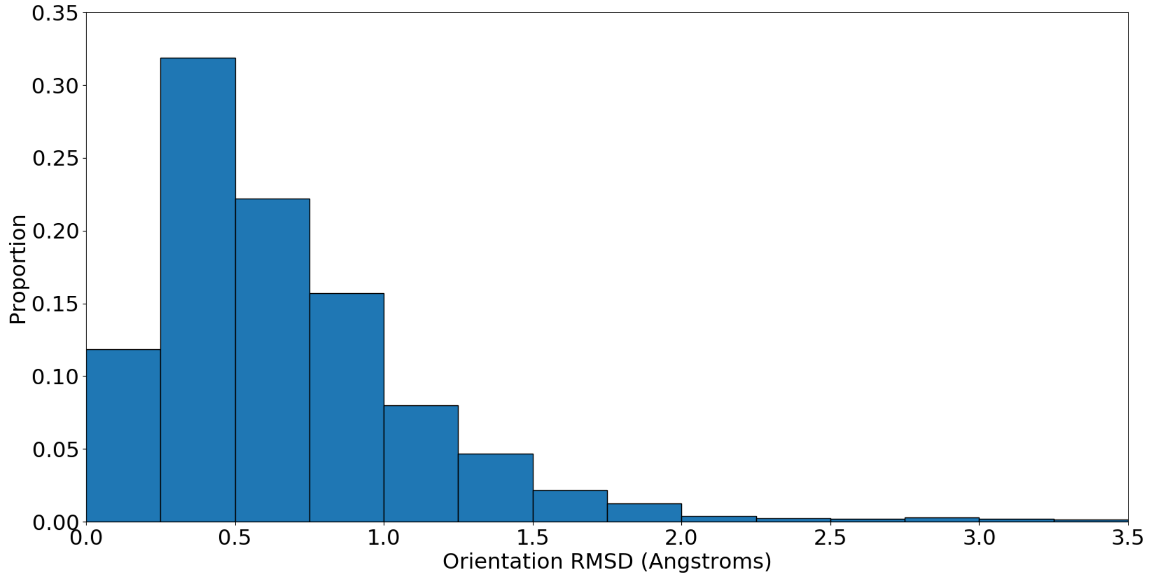


**Fig. 2.** The distribution of orientation RMSDs observed between Fvs of identical heavy and light chain sequence. The vast majority (92%) have orientation RMSDs below 1.5Å.

DRAFT

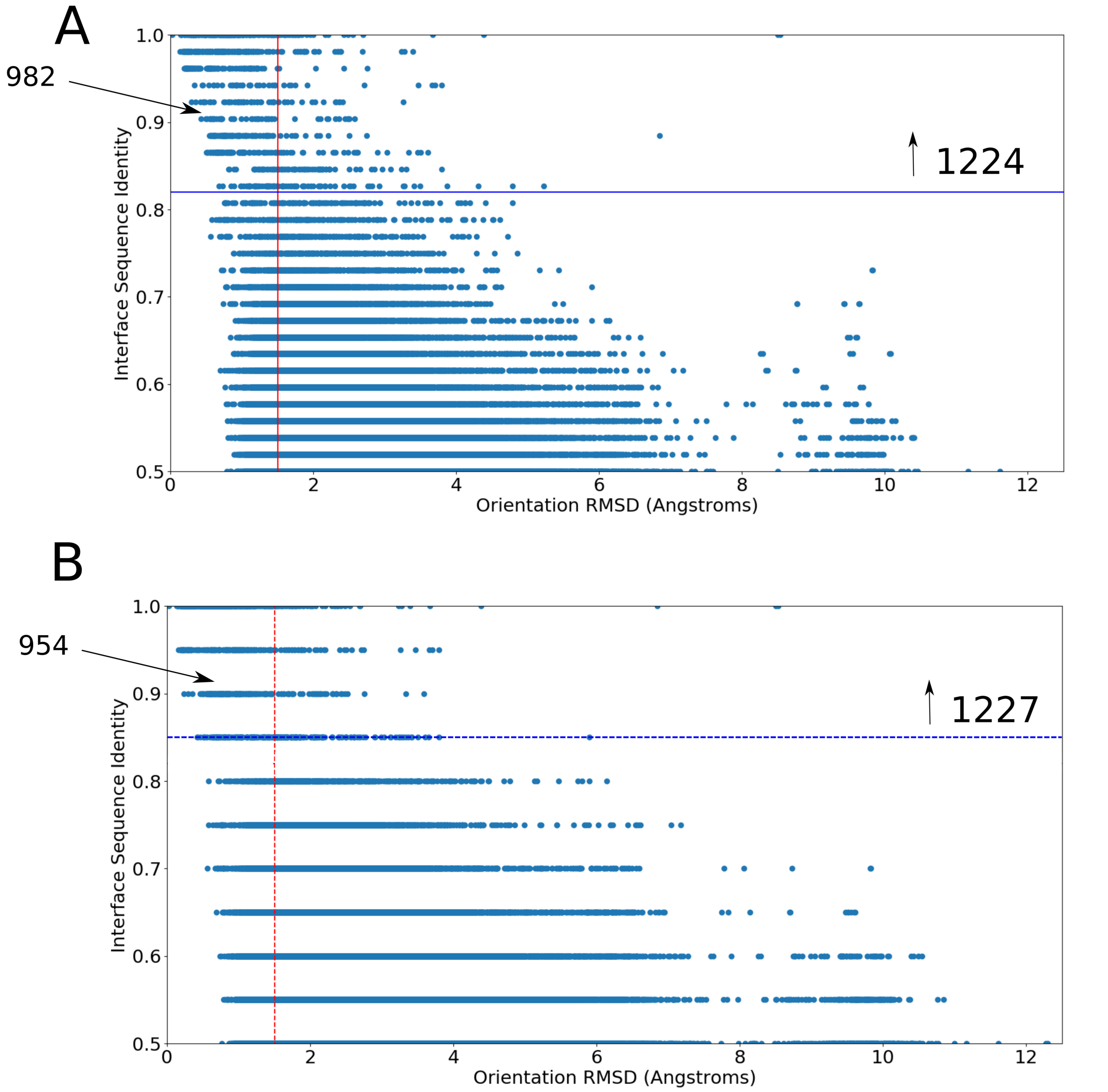


**Fig. 3.** Graphs showing the orientation RMSD observed at each interface sequence identity value for (A) all 52 interface residues and (b) the 20 most important interface residues. The thresholds for (A) are set at 1.5Å and 82% sequence identity, while for (B) are set at 1.5Å and 85% sequence identity. The proportions above the sequence identity threshold and within 1.5Å orientation RMSD are 80.2% (982/1224) and 77.8% (954/1227) respectively.

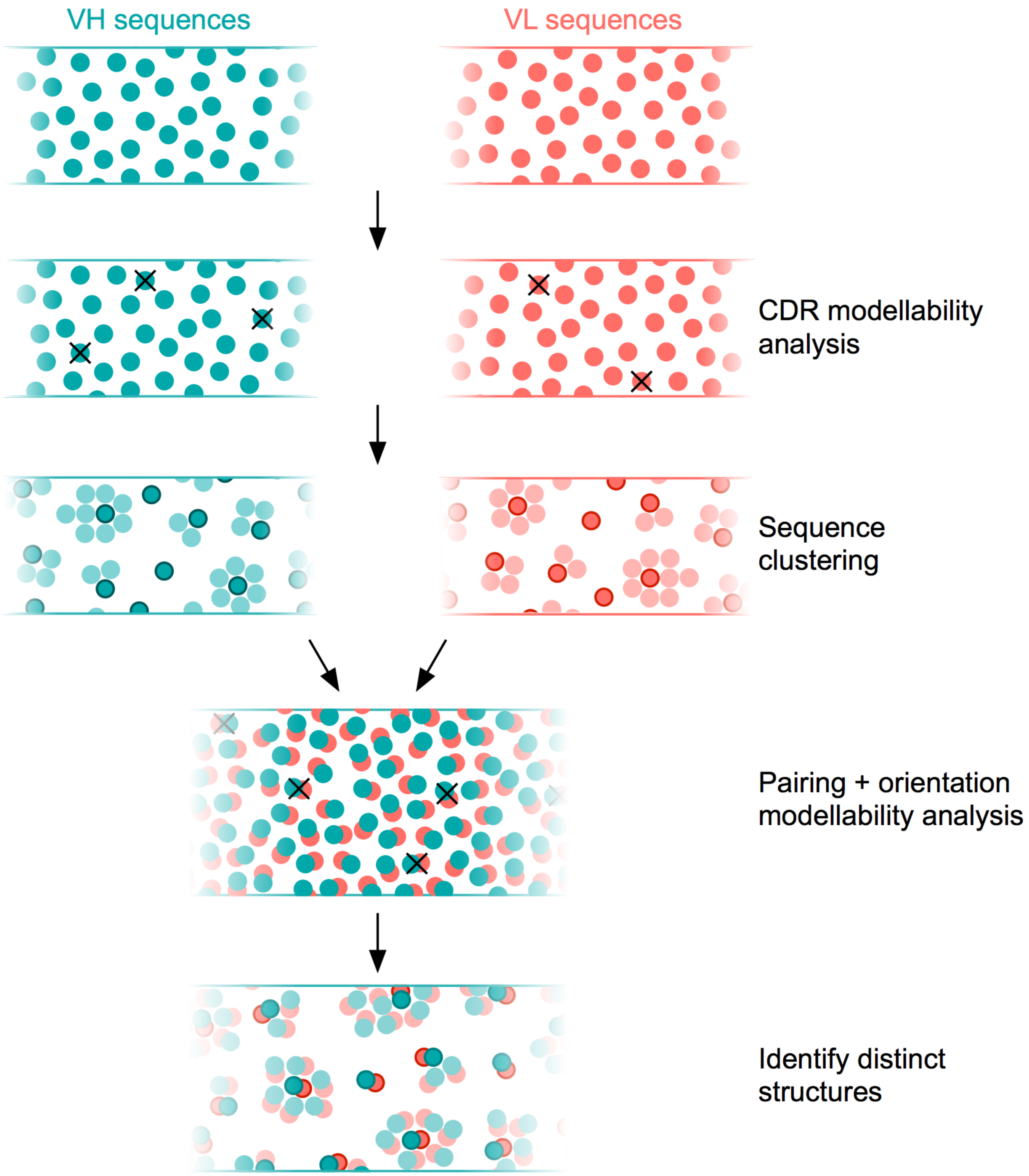


**Fig. 4.** A schematic illustrating our repertoire structural profiling algorithm. Heavy (VH) and light (VL) chain sequences from a repertoire snapshot are first analysed separately for their FREAD (1, 2) modellability (unmodellable chains are crossed out). They are then clustered by sequence identity using CD-HIT (6) (90% threshold) for computational tractability. All VH and VL cluster centre chains are subsequently paired, and VH-VL orientations that cannot reliably modelled are removed (again shown by crosses). Finally, predicted modellable Fvs with identical combinations of CDR lengths are structurally clustered to identify 'distinct structures'.

| Interface Residues | |
|---|---|
| Heavy Chain | Light Chain |
| **44** | 36 |
| **47** | 37 |
| **48** | 38 |
| 49 | 40 |
| 50 | 42 |
| **52** | 44 |
| 55 | 47 |
| 57 | 48 |
| 66 | 49 |
| 68 | **50** |
| 69 | 52 |
| 70 | 55 |
| **101** | **56** |
| 103 | 66 |
| **107** | 68 |
| 108 | **69** |
| **109** | **101** |
| 113 | **103** |
| **114** | 105 |
| 115 | 107 |
| **116** | 108 |
| **117** | **109** |
| 118 | 114 |
| 119 | **115** |
| **120** | **116** |
| | 118 |
| | **120** |

DRAFT

**Table 1.** The 52 heavy and light chain residues tending to lie in the heavy-light chain interface. Residue numbers in bold were determined to be amongst the five most important in the Random Forest regression model when predicting the six different ABangle parameters.

| Dataset | All VH | All VL | Modellable VH [90% SIC] | Modellable VL [90% SIC] | Predicted Modellable Fvs | Distinct Structures |
|---|---|---|---|---|---|---|
| 1 (S64) | 177,603 | 123,934 | 10,087 | 6,779 | 6,420,211 | 209,394 |
| 2 (S57) | 169,805 | 118,020 | 9,860 | 7,922 | 7,225,630 | 201,039 |
| 3 (S5) | 159,544 | 139,845 | 8,999 | 8,526 | 6,827,419 | 200,708 |
| 4 (S56) | 162,446 | 136,874 | 9,309 | 7,168 | 6,628,683 | 195,061 |
| 5 (S83) | 152,299 | 112,733 | 9,048 | 8,076 | 6,170,373 | 193,384 |
| 6 (S67) | 173,722 | 120,237 | 9,349 | 6,424 | 5,544,952 | 193,061 |
| 7 (S84) | 164,017 | 138,874 | 8,702 | 8,232 | 5,634,598 | 191,617 |
| 8 (S76) | 148,180 | 126,713 | 8,778 | 7,047 | 5,856,150 | 191,162 |
| 9 (S54) | 121,993 | 133,921 | 7,581 | 9,066 | 5,074,822 | 181,290 |
| 10 (S89) | 152,710 | 144,340 | 8,923 | 9,293 | 5,414,820 | 177,829 |
| 11 (S13) | 127,321 | 134,485 | 7,276 | 8,654 | 5,314,377 | 174,105 |
| 12 (S93) | 110,676 | 110,904 | 7,260 | 7,528 | 4,799,497 | 173,006 |
| 13 (S87) | 120,424 | 104,440 | 7,569 | 6,145 | 4,043,317 | 172,200 |
| 14 (S86) | 156,096 | 132,580 | 8,411 | 7,475 | 5,130,237 | 171,940 |
| 15 (S10) | 109,552 | 134,816 | 6,816 | 8,351 | 5,152,331 | 167,153 |
| 16 (S50) | 157,428 | 109,437 | 8,614 | 5,533 | 4,556,841 | 162,663 |
| 17 (S75) | 105,099 | 119,470 | 6,576 | 7,711 | 4,174,078 | 156,510 |
| 18 (S8) | 150,763 | 112,479 | 8,241 | 6,604 | 4,305,148 | 156,044 |
| 19 (S37) | 137,951 | 103,825 | 7,815 | 5,983 | 3,565,942 | 156,034 |
| 20 (S59) | 111,842 | 125,621 | 6,598 | 8,117 | 4,807,933 | 150,425 |
| 21 (S22) | 113,023 | 145,227 | 6,220 | 8,781 | 4,108,518 | 149,575 |
| 22 (S77) | 118,326 | 114,384 | 7,407 | 7,195 | 3,821,870 | 148,950 |
| 23 (S58) | 114,817 | 101,696 | 7,663 | 7,088 | 4,388,190 | 148,753 |
| 24 (S30) | 112,518 | 107,459 | 6,383 | 6,842 | 3,667,996 | 148,691 |
| 25 (S27) | 125,328 | 122,194 | 7,347 | 7,305 | 3,989,334 | 147,654 |
| 26 (S74) | 119,371 | 104,430 | 6,375 | 5,753 | 2,961,937 | 146,049 |
| 27 (S12) | 117,699 | 127,682 | 6,762 | 6,860 | 3,388,406 | 144,916 |
| 28 (S19) | 111,012 | 102,905 | 6,595 | 6,064 | 3,471,725 | 143,027 |
| 29 (S52) | 109,826 | 113,067 | 6,450 | 5,760 | 3,397,333 | 142,110 |
| 30 (S60) | 112,802 | 108,429 | 6,513 | 5,790 | 3,310,781 | 140,525 |
| 31 (S11) | 102,847 | 114,304 | 5,610 | 6,873 | 3,495,280 | 139,094 |
| 32 (S51) | 117,156 | 122,970 | 6,187 | 6,652 | 3,828,676 | 139,024 |
| 33 (S25) | 108,425 | 101,548 | 6,159 | 6,125 | 3,176,321 | 137,503 |
| 34 (S34) | 114,748 | 174,784 | 6,195 | 11,937 | 6,404,104 | 136,973 |
| 35 (S29) | 100,214 | 143,027 | 5,369 | 9,501 | 3,819,414 | 136,811 |
| 36 (S72) | 118,995 | 111,611 | 6,360 | 6,152 | 2,627,255 | 136,161 |
| 37 (S91) | 114,196 | 118,422 | 6,175 | 8,304 | 4,256,118 | 132,851 |
| 38 (S65) | 134,646 | 124,116 | 7,008 | 8,584 | 3,446,622 | 108,021 |
| 39 (S95) | 118,576 | 162,377 | 5,412 | 11,748 | 5,901,443 | 91,855 |
| 40 (S17) | 102,405 | 111,669 | 5,310 | 7,945 | 2,690,081 | 91,229 |
| 41 (S4) | 100,689 | 128,986 | 4,688 | 1,761 | 745,977 | 78,588 |
| *Overall* | | | | | *183,544,740* | |

DRAFT

**Table 2.** Structurally profiling the baseline repertoire snapshots of 41 unrelated individuals (7). In order, the columns show: the dataset label, the number of VH and VL reads within each snapshot, the number of FREAD-modellable VH and VL reads (once clustered at 90% sequence identity), the number of predicted modellable Fvs resulting from these VH-VL pairings, and the number of distinct structures (cluster centres) identified in each dataset. SIC = Sequence Identity Clustered.

| # of Repertoires (Dataset Added) | Modellable Fvs Added | Cumulative Public & Private Distinct Structures | Public Distinct Structures (Overall % Public) |
|---|---|---|---|
| 1 (S64) | 6,420,211 | 209,394 | 209,394 |
| 2 (+S57) | 7,225,630 | 340,915 | 100,824 (29.57) |
| 3 (+S5) | 6,827,419 | 445,045 | 71,743 (16.12) |
| 4 (+S56) | 6,628,683 | 527,668 | 58,043 (11.00) |
| 5 (+S83) | 6,170,373 | 604,124 | 48,703 (8.06) |
| 6 (+S67) | 5,544,952 | 670,833 | 42,277 (6.30) |
| 7 (+S84) | 5,624,598 | 734,374 | 37,151 (5.06) |
| 8 (+S76) | 5,856,150 | 793,831 | 33,572 (4.23) |
| 9 (+S54) | 5,074,822 | 846,670 | 30,474 (3.60) |
| 10 (+S89) | 5,414,820 | 896,328 | 27,389 (3.06) |
| 11 (+S13) | 5,314,377 | 940,957 | 25,621 (2.72) |
| 12 (+S93) | 4,799,497 | 980,905 | 24,015 (2.45) |
| 13 (+S87) | 4,043,317 | 1,023,105 | 22,052 (2.16) |
| 14 (+S86) | 5,130,237 | 1,061,003 | 20,867 (1.97) |
| 15 (+S10) | 5,152,331 | 1,100,394 | 19,468 (1.77) |
| 16 (+S50) | 4,556,841 | 1,130,974 | 18,421 (1.63) |
| 17 (+S75) | 4,174,078 | 1,161,111 | 17,157 (1.48) |
| 18 (+S8) | 4,305,148 | 1,188,715 | 16,071 (1.35) |
| 19 (+S37) | 3,565,942 | 1,218,071 | 15,302 (1.26) |
| 20 (+S59) | 4,807,933 | 1,243,044 | 14,669 (1.18) |
| 21 (+S22) | 4,108,518 | 1,269,972 | 13,992 (1.11) |
| 22 (+S77) | 3,821,870 | 1,294,338 | 13,380 (1.03) |
| 23 (+S58) | 4,388,190 | 1,316,084 | 12,953 (0.98) |
| 24 (+S30) | 3,667,996 | 1,342,141 | 12,542 (0.93) |
| 25 (+S27) | 3,989,334 | 1,365,687 | 12,017 (0.88) |
| 26 (+S74) | 2,961,937 | 1,387,177 | 11,482 (0.83) |
| 27 (+S12) | 3,388,406 | 1,406,918 | 11,050 (0.79) |
| 28 (+S19) | 3,471,725 | 1,426,572 | 10,732 (0.75) |
| 29 (+S52) | 3,397,333 | 1,446,838 | 10,319 (0.71) |
| 30 (+S60) | 3,310,781 | 1,466,071 | 10,078 (0.69) |
| 31 (+S11) | 3,495,280 | 1,486,113 | 9,714 (0.65) |
| 32 (+S51) | 3,828,676 | 1,504,633 | 9,478 (0.63) |
| 33 (+S25) | 3,176,321 | 1,522,848 | 9,141 (0.60) |
| 34 (+S34) | 6,404,104 | 1,544,438 | 8,615 (0.56) |
| 35 (+S29) | 3,819,414 | 1,564,003 | 8,226 (0.53) |
| 36 (+S72) | 2,627,255 | 1,581,699 | 7,966 (0.50) |
| 37 (+S91) | 4,256,118 | 1,598,785 | 7,818 (0.49) |
| 38 (+S65) | 3,446,622 | 1,614,661 | 6,891 (0.43) |
| 39 (+S95) | 5,901,443 | 1,629,262 | 5,935 (0.41) |
| 40 (+S17) | 2,690,081 | 1,642,531 | 5,110 (0.31) |
| 41 (+S4) | 745,977 | 1,650,922 | 4,573 (0.28) |

DRAFT

**Table 3.** Evaluating the number of public distinct structures seen across multiple baseline repertoire snapshots. In order, the columns show: the number of repertoires compared (in brackets the identifier of the last dataset added), the number of predicted modellable Fvs added by the last dataset, the number of distinct structures added by the last dataset, the (cumulative) number of public and private distinct structures across all compared repertoires, and the number of proportion of these structures that are public. The sharp drop-off in the proportion of public structures in the final four repertoire snapshots can be rationalised by their substantially lower internal structural diversity (see Table 2).

| # of Repertoires (Dataset Added) | Cumulative Modellable Fvs | Cumulative Distinct Structures | Expected Cumulative Distinct Structures |
|---|---|---|---|
| 1 (S64) | 6,420,211 | 209,394 | 209,394 |
| 2 (+S57) | 13,645,841 | 340,915 | 445,057 |
| 3 (+S5) | 20,473,260 | 445,045 | 667,732 |
| 4 (+S56) | 27,101,943 | 527,668 | 883,925 |
| 5 (+S83) | 33,272,316 | 604,124 | 1,085,170 |
| 6 (+S67) | 38,817,268 | 670,833 | 1,266,018 |
| 7 (+S84) | 44,451,866 | 734,374 | 1,449,463 |
| 8 (+S76) | 50,308,016 | 793,831 | 1,640,461 |
| 9 (+S54) | 55,382,838 | 846,670 | 1,805,975 |
| 10 (+S89) | 60,797,658 | 896,328 | 1,982,578 |
| 11 (+S13) | 66,112,035 | 940,957 | 2,156,232 |
| 12 (+S93) | 70,911,532 | 980,905 | 2,312,767 |
| 13 (+S87) | 74,954,849 | 1,023,105 | 2,444,639 |
| 14 (+S86) | 80,085,086 | 1,061,003 | 2,611,960 |
| 15 (+S10) | 85,237,417 | 1,100,394 | 2,780,003 |
| 16 (+S50) | 89,794,258 | 1,130,974 | 2,928,623 |
| 17 (+S75) | 93,968,336 | 1,161,111 | 3,064,760 |
| 18 (+S8) | 98,273,484 | 1,188,715 | 3,205,172 |
| 19 (+S37) | 101,839,426 | 1,218,071 | 3,321,474 |
| 20 (+S59) | 106,647,359 | 1,243,044 | 3,478,284 |
| 21 (+S22) | 110,755,877 | 1,269,972 | 3,612,283 |
| 22 (+S77) | 114,577,747 | 1,294,338 | 3,736,932 |
| 23 (+S58) | 118,965,937 | 1,316,084 | 3,880,052 |
| 24 (+S30) | 122,633,933 | 1,342,141 | 3,999,683 |
| 25 (+S27) | 126,623,267 | 1,365,687 | 4,129,795 |
| 26 (+S74) | 129,585,204 | 1,387,177 | 4,226,398 |
| 27 (+S12) | 132,973,610 | 1,406,918 | 4,336,910 |
| 28 (+S19) | 136,445,335 | 1,426,572 | 4,450,139 |
| 29 (+S52) | 139,842,668 | 1,446,838 | 4,560,943 |
| 30 (+S60) | 143,153,449 | 1,466,071 | 4,668,923 |
| 31 (+S11) | 146,648,729 | 1,486,113 | 4,782,921 |
| 32 (+S51) | 150,477,405 | 1,504,633 | 4,907,793 |
| 33 (+S25) | 153,653,726 | 1,522,848 | 5,011,389 |
| 34 (+S34) | 160,057,830 | 1,544,438 | 5,220,257 |
| 35 (+S29) | 163,877,244 | 1,564,003 | 5,344,827 |
| 36 (+S72) | 166,504,499 | 1,581,699 | 5,430,514 |
| 37 (+S91) | 170,760,617 | 1,598,785 | 5,569,326 |
| 38 (+S65) | 174,207,239 | 1,614,661 | 5,681,737 |
| 39 (+S95) | 180,108,682 | 1,629,262 | 5,874,212 |
| 40 (+S17) | 182,798,763 | 1,642,531 | 5,961,948 |
| 41 (+S4) | 183,544,740 | 1,650,922 | 5,986,278 |

**Table 4.** Tracking the total number of public and private distinct structures seen across multiple baseline repertoire snapshots. In order, the columns show: the number of repertoires compared (in brackets the identifier of the last dataset added), the cumulative number of predicted modellable Fvs, the number of public and private distinct structures seen across all compared repertoires, and the expected number of cumulative public and private distinct structures if new distinct structures were observed at the same rate per modellable Fv as seen in S64.

DRAFT

| PB AML Structure | Therapeutic | PDB ID | Fv RMSD (Å) | CDR Lengths (H1-3, L1-3) | Antigen Target |
|---|---|---|---|---|---|
| H101+L64549 | Ofatumumab | 3giz (HL) | 0.148 | 13-10-15-11-8-9 | CD20 |
| H11835+L101012 | Durvalumab | 5xj4 (HL) | 0.162 | 13-10-14-12-8-9 | CD274 |
| H19709+L100051 | Tanezumab | 4edw (HL) | 0.181 | 13-9-15-11-8-9 | NGFB |
| H35853+L102278 | Tremelimumab | 5ggu (HL) | 0.255 | 13-10-18-11-8-9 | CD152 |
| H11488+L100048 | Olokizumab | 5tru (HL) | 0.278 | 13-12-11-11-8-9 | IL6 |
| H10992+L14321 | Tezepelumab | 5j13 (CB) | 0.481 | 13-10-15-11-8-11 | TSLP |
| H13677+L17885 | Avelumab | 4nki (HL) | 0.525 | 13-10-13-14-8-10 | CD274 |
| H14012+L14649 | Ustekinumab | 3hmw (HL) | 0.638 | 13-10-12-11-8-9 | IL12B |
| H38817+L68369 | Aducanumab | 6cnr (HL) | 0.681 | 13-10-17-11-8-9 | APP |
| H26854+L108275 | Imalumab | 6foe (HL) | 0.730 | 13-10-11-11-8-9 | MIF |
| H30607+L11664 | Quilizumab | 3hr5 (HL) | 0.730 | 13-10-10-16-8-9 | IGHE |

**Table 5.** The eleven clinical-stage therapeutic antibodies with a solved crystal structure within 0.75Å variable domain (Fv) root-mean-squared deviation (RMSD) of an antibody model structure from the Public Baseline Antibody Model Library (PB AML). The first column records the Fv identifier for the geometrically closest AML model to each of the eleven therapeutics listed in column 2. Column 3 provides the Protein Data Bank (PDB) identifier for each chosen therapeutic structure (chain identifiers in brackets). The corresponding RMSD is provided in column 4; all RMSD comparisons were made between AML structures and therapeutics with an identical combination of CDR lengths. This combination of North-defined (8) CDR lengths is then listed in the order H1-H2-H3-L1-L2-L3. Finally, the target for each therapeutic antibody is recorded. PDB = Protein Data Bank; VH = variable heavy chain; VL = variable light chain; Fv = Fragment variable region; RMSD = root-mean-squared deviation; CDR = Complementarity-Determining Region.

Antigens: CD - Cluster of Differentiation protein, NGFB - Nerve Growth Factor B, IL - interleukin, TSLP - Thymic Stromal Lymphopoietin, APP - Amyloid Precursor Protein, MIF - Macrophage Migration Inhibitory Factor, IGHE - Immunoglobulin Heavy Constant Epsilon.

DRAFT

| Repertoire | All VH | All VL | Modellable VH [90% SIC] | Modellable VL [90% SIC] | Predicted Modellable Fvs | Distinct Structures |
|---|---|---|---|---|---|---|
| V1 Before | 241,630 | 679,472 | 13,230 | 93,422 | 33,376,943 | 471,650 |
| V2 Before | 235,022 | 754,179 | 12,135 | 105,026 | 47,884,336 | 579,961 |
| V3 Before | 307,067 | 825,587 | 16,124 | 113,239 | 77,186,291 | 772,844 |
| V1 After | 392,661 | 1,052,510 | 21,624 | 155,412 | 104,736,939 | 852,328 |
| V2 After | 331,573 | 1,173,628 | 16,047 | 158,439 | 91,023,995 | 809,341 |
| V3 After | 368,961 | 1,087,670 | 18,029 | 299,908 | 105,824,532 | 843,341 |

**Table 6.** Structurally profiling the 'Before Vaccination' (Before) and 'After Vaccination' (After) repertoire snapshots of three unrelated individuals (V1, V2, and V3) (9) . In order, the columns show: the dataset label, the number of VH and VL reads within each snapshot, the number of FREAD-modellable VH and VL reads (once clustered at 90% sequence identity), the number of predicted-modellable Fvs resulting from these VH-VL pairings, and the number of distinct structures (cluster centres) identified through greedy structural clustering. SIC = Sequence Identity Clustered.

DRAFT

### Datasets

The following datasets are hosted at *http://opig.stats.ox.ac.uk/resources*:

1. *Public_Baseline_AML.tar.gz*

The 23,700 ABodyBuilder variable domain homology models representing the Public Baseline Antibody Model Library.

2. *Public_Response_AML.tar.gz*

The 74,181 ABodyBuilder variable domain homology models representing the Public Response Antibody Model Library.


DRAFT
